# Young and Undamaged rMSA Improves the Longevity of Mice

**DOI:** 10.1101/2021.02.21.432135

**Authors:** Jiaze Tang, Anji Ju, Boya Li, Shaosen Zhang, Yuanchao Gong, Boyuan Ma, Yi Jiang, Hongyi Liu, Yan Fu, Yongzhang Luo

**Affiliations:** The National Engineering Laboratory for Anti-Tumor Protein Therapeutics, Tsinghua University, Beijing, China; Beijing Key Laboratory for Protein Therapeutics, Tsinghua University, Beijing, China; Cancer Biology Laboratory, School of Life Sciences, Tsinghua University, Beijing, China

**Keywords:** rMSA, longevity, free thiol, advanced glycation end-product, carbonyl, homocysteine.

## Abstract

Improvement of longevity is an eternal dream of human beings. Here we report that a single protein recombinant mouse serum albumin (rMSA) improved the lifespan and healthspan of C57BL/6N mice. The median lifespan extensions were 17.6% for female and 20.3% for male, respectively. The grip strength of rMSA-treated female and male mice increased by 29.6% and 17.4%, respectively. Meanwhile, the percentage of successful escape increased 23.0% in rMSA-treated male mice using the Barnes Maze test. The rMSA used in this study is young and almost undamaged. We define the concept “young and undamaged” to any protein without any unnecessary modifications by four parameters: intact free thiol (if any), no advanced glycation end-product, no carbonylation, and no homocysteinylation. Here “young and undamaged” rMSA is much younger and less damaged than the endogenous serum albumin from young mice at 1.5 months of age. We predict that young and undamaged proteins altogether can further improve the longevity.

---

Longevity is an eternal pursuit of human beings. Tales of passionate seeking for immortality ran through the whole human history. Ludwig et al reported the extended lifespan of older rats by younger rats in the parabiosis model for the first time in 1972 [1]. Egerman group and Villeda group respectively found that the muscle strength and cognitive ability of old mice were improved after the parabiosis surgery with young mice [2, 3], which suggest that the “mystery” of aging may exist in blood proteins. It is believed that aging is at least partially caused by the continuous accumulation of damages or unnecessary modifications of proteins [4, 5, 6], including free thiol oxidation, carbonylation, advanced glycation end-product (AGE) formation, and homocysteinylation [7, 8, 9, 10].

Human serum albumin (HSA, UniProtKB P02768) is the most abundant protein in blood plasma with a serum half-life of about 21 days [11]. Damages or unnecessary modifications of HSA are related to many pathological conditions and increase with age. Firstly, the single free thiol in Cys-34 residue of HSA has been proposed to account for approximately 80% of the total free thiols in plasma [12, 13], whose oxidation is intimately linked with aging and age-related diseases [14, 15, 16]. Secondly, in oxidative environments, carbonyls are also formed especially on the side chains of Pro, Arg, Lys and Thr residues in proteins [17, 18]. Elevated carbonyl levels in HSA have been found to be related to aging and varieties of diseases [19, 20, 21]. Thirdly, the AGE accumulation of HSA is another important factor found to be involved in aging [9, 22]. It is widely reported that AGE formation impairs normal functions of albumin and can induce inflammatory responses, which is connected with aging and the progression of serious diseases [22, 23]. Fourthly, it has been widely reported that homocysteine (Hcy) increases with age and is associated with age-related degenerative disorders [10, 24, 25, 26]. HSA is a major target for homocysteinylation, thus it can efficiently protect other proteins from the toxicity of Hcy [27, 28, 29].

Therefore, treatment of freshly prepared recombinant serum albumin with no damages or unnecessary modifications is most likely to extend lifespan and healthspan. Here we report that young and undamaged recombinant mouse serum albumin (rMSA)-treated groups in natural aging mouse model obtained significantly extended lifespan with increased skeletal muscle strength and cognitive ability compared with saline-treated groups.

## Materials and Methods

### Mice and drug treatments

C57BL/6N mice were purchased from Beijing Vital River Laboratory Animal Technology Co., Ltd. (a distributor of Charles River Laboratories in China). The mice transport stress syndrome was carefully avoided during the transportation to the Laboratory Animal Research Center, Tsinghua University (THU-LARC). All mice were quarantined for one month to guarantee the adaptation to the new environment and carried out quality inspection. Animals were kept in a pathogen-free barrier environment with a 12-h dark-light circle. Room temperature was maintained at 23 °C. After arrival, mice were fed with irradiation-sterilized JAX-standard breeder chow (SHOOBREE^®^, Xietong Pharmaceutical Bio-technology Co., Ltd., 1010058) and sterilized water during the entire study.

12-month-old middle aged mice were divided into rMSA- or saline-treated group randomly. More than one kilogram correctly refolded rMSA was kindly supplied by Shenzhen Protgen, Ltd. The quality of GMP-grade rMSA, expressed by *pichia pastoris*, was strictly controlled to ensure that the purity is greater than 99%. Most importantly, host cell proteins (HCPs) were less than 1 μg/g rMSA by ELISA, which means our rMSA is almost free of HCPs.

125 mg/mL of rMSA dissolved in saline was *i.v.* injected slowly. Mice were weighed before each injection to calculate the dosage, with saline served as the negative control. Mice were injected with 1.5 mg rMSA per gram of mouse body weight and isometric saline every 3 weeks as indicated. All animal studies were approved by the Institutional Animal Care and Use Committee of Tsinghua University (Beijing, China).

### Protein levels determination

To determine the blood biochemical parameters, blood samples were collected from mouse orbital sinus after Avertin^®^ (Tribromoethanol, Sigma-Aldrich, T48402) intraperitoneal injection (400 mg/kg) for anesthesia. Serum samples were collected after centrifugation at 1,000 × g for 20 min at 4 °C. To collect plasma samples, heparin sodium salt is added to the fresh blood samples (20 units/mL blood, Sigma-Aldrich, H3149) to prevent blood clotting followed by centrifugation at 1,000 × g for 30 min at 4 °C. Major blood biochemical parameters of serum samples were determined with an automatic biochemistry analyzer (Olympus AU 400).

To determine the expression level of albumin, mice were euthanized using carbon dioxide after anesthesia. Liver tissue samples were quickly removed and homogenized. The total RNA from the homogenate was isolated using TRIzol Reagent (Invitrogen, 15596026) and converted into cDNA using the First Strand cDNA Synthesis Kit (Fermentas, K1622). Quantitative RT-PCR (qRT-PCR) was performed using the TransStart^®^ Top Green qPCR SuperMix (TransGen Biotech Co., 1). Relative quantitation was analyzed using the ΔΔ Ct method. Glyceraldehyde 3-phosphate dehydrogenase (GAPDH) was used as an internal control. Independent experiments were repeated in triplicates. The following primers were used: *Alb* forward 5’-TGCTTTTCCAGGGGTGTGTT, reverse 5’-TTACTTCCTGCACTAATTTGGCA; *Gapdh* forward 5’-GTTGTCTCCTGCGACTTCA, reverse 5’-GGTGGTCCAGG GTTTCTTA.

### Grip strength test

The grip strength was measured using a grip strength meter (Yiyan Co. Ltd., YLS-13A). Mice were allowed to hold on to a metal grid and were gently pulled backwards by the tail at a constant speed until the mice could no longer hold the grid. Each mouse was given five trials, and the average value was used to represent the grip strength of an individual mouse. The experiments were carried out in a randomized double-blind procedure.

### Barnes maze assay

Male mice treated with rMSA or isometric saline for 8 months were subjected to the Barnes maze assay to evaluate spatial memory function. For the Barnes maze assay, mice were trained to find a hole that connected to a black escape box, which was positioned around the circumference of a circular platform (Shanghai XinRuan, XR-XB108). The circular platform was 91 cm diameter and 0.4 cm thick, with 20 evenly distributed 5 cm diameter holes around the edge, with two overhead lights served as an aversive stimulus. Each trial was recorded by a video camera installed over the platform. Procedures were similar as described by Rosenfeld et al with modifications [30]. The results were analyzed by Super Maze software. The experiments were carried out in a randomized double-blind procedure.

### Albumin purification

Serum samples of indicated groups were diluted with 20 mM Tris buffer containing 0.15 M NaCl at pH 7.8 before applying to a pre-equilibrated Blue BestaroseTMFF column (Bestchrom), followed by 3-bed volumes wash of nonspecific binding proteins. Mouse albumin was eluted by elution buffer (0.2 M NaSCN, pH 8.0), then dialyzed against PBS and concentrated by Amicon® ultra centrifugal filters with Ultracel-30 regenerated cellulose membrane (MerckMillipore, UFC803008) at 4°C. Protein concentrations were determined by the Pierce™ BCA Protein Assay Kit according to manufacturer’s instructions (Thermo Scientific, 23227). Samples were analyzed on a Quadrupole-Time of Flight (Q-TOF) mass spectrometer (Waters, SYNAPT G2-Si) instrument optimized for high-mass protein molecular weight analysis.

### Immunofluorescence assay

Frozen sections of mice which were dissected from mice, fixed with cold acetone. Then these samples were blocked with 10% goat serum and stained with primary antibodies overnight at 4°C followed by the appropriate secondary fluorescently labeled antibodies at 4 °C overnight. Slides were stained with FITC-conjugated secondary antibodies, and nuclei were stained by 4′,6-diamidino-2-phenylindole (DAPI). Fluorescence imaging was performed on Nikon A1 laser scanning confocal microscope and was analyzed with NIS-Elements Software (Nikon) and ImageJ software.

The following antibodies were used: mouse monoclonal antibody against phosphorylated microtubule-associated protein tau (p-tau, UniProtKB P10637. Thermo Fisher Scientific, MN1020), mouse monoclonal antibody against slow myosin heavy chain I (MYH1, UniProtKB Q5SX40. Sigma-Aldrich, M8421), rabbit monoclonal antibody against α-smooth muscle actin (α-SMA, UniProtKB P62737. Cell Signaling Technology, 19245), FITC-conjugated goat polyclonal antibody against mouse IgG (H+L) (Abcam, ab6785), and FITC-conjugated goat polyclonal antibody against rabbit IgG (H+L) (Abcam, ab97050).

### Masson’s trichrome staining

Paraformaldehyde-fixed, paraffin-embedded tissue sections from mice were deparaffinized and rehydrated. Then sections were stained with the Masson’s Trichrome Stain Kit (KeyGEN BioTECH, KGMST-8004). Nuclei stain black, cytoplasm and muscle fibers stain red, and collagen displays blue coloration.

### Toluidine Blue O staining

Paraformaldehyde-fixed, paraffin-embedded tissue sections from mice were deparaffinized and rehydrated. Then sections were stained with the Toluidine Blue O reagent according to manufacturer’s instructions (Solarbio, G3668).

### Immunohistochemical assay

The rehydrated sections were rinsed three times with PBS and the endogenous peroxidase was blocked with 3% H_2_O_2_. Then the samples were blocked with 10% goat serum and incubated with primary antibodies overnight at 4°C followed by the appropriate secondary HRP-conjugated antibodies at 4°C overnight. Slides were stained with newly prepared DAB substrate and nuclei were stained by hematoxylin. The immunohistochemical staining intensity was quantified with ImageJ software.

The following antibodies were used: rabbit monoclonal antibody against collagen I (COL1A1, UniProtKB P11087. Cell Signaling Technology, 91144), rabbit polyclonal antibody against desmin (UniProtKB P31001. Thermo Fisher Scientific, PA5-16705), rabbit monoclonal antibody against α-SMA (Cell Signaling Technology, 19245), and HRP-conjugated goat polyclonal antibody against rabbit IgG (H+L) (Abcam, ab205718).

### Aging-related parameters determination

The Ellman’s method was used to determine the content of free thiols [31]. Mouse serum albumin (MSA, UniProtKB P07724) and rMSA were mixed with equal volumes of 5, 5’-Dithiobis-(2-nitrobenzoic acid) (DTNB) reagent, respectively. The volume and concentration of DTNB used in this study were 100 μL and 2 mM, respectively. 800 μL Tris buffer (1 M) was added to make the volume of the reaction system reach 1000 μL. Samples were kept at room temperature for 30 min. The fluorescence absorbance was measured at 412 nm. Carbonyls in protein samples were quantified using the Protein Carbonyl Content Assay Kit (Abcam, ab126287) according to the manual. Hcy concentrations were measured by the enzyme-linked immunosorbent assay (ELISA) according to manufacturer’s instructions (MEIMIAN, 1213). Concentrations of AGE were measured with an ELISA kit according to manufacturer’s instructions (CLOUD-CLONE Co., CEB353Ge).

### Statistical analysis

The Kaplan–Meier method was used for survival analysis and the survival curves were compared by using the log-rank (Mantel-Cox) test. The variance across samples was analyzed using Kolmogorov-Smirnov (K-S) test and Levene’s test, followed by 2-tailed unpaired Student *t*-test, where *p* < 0.05 is considered significant. Statistical analysis and diagramming were carried out by the Graphpad Prism 6.01 software unless otherwise noted.

## Results

### rMSA treatment increased the longevity in mice

In order to verify whether rMSA treatment can extend the lifespan of mice, 12-month-old middle aged C57BL/6N mice were chosen as natural aging models, and were *i.v.* injected with 1.5 mg rMSA per gram of body weight or isometric saline every 3 weeks until natural death. The lifespans of rMSA-treated mice were improved significantly (Fig. 1A), wherein 17.6% for females (3.4 months increased, *p* = 0.0164, Fig. 1B) and 20.3% for males (3.9 months increased, *p* = 0.0342, Fig. 1C). Changes in the appearance of both sexes were observed when the median lifespan was reached. Interestingly, mice treated with rMSA had glossier and thicker fur than saline-treated mice (Fig. 1D).

**Figure 1.**
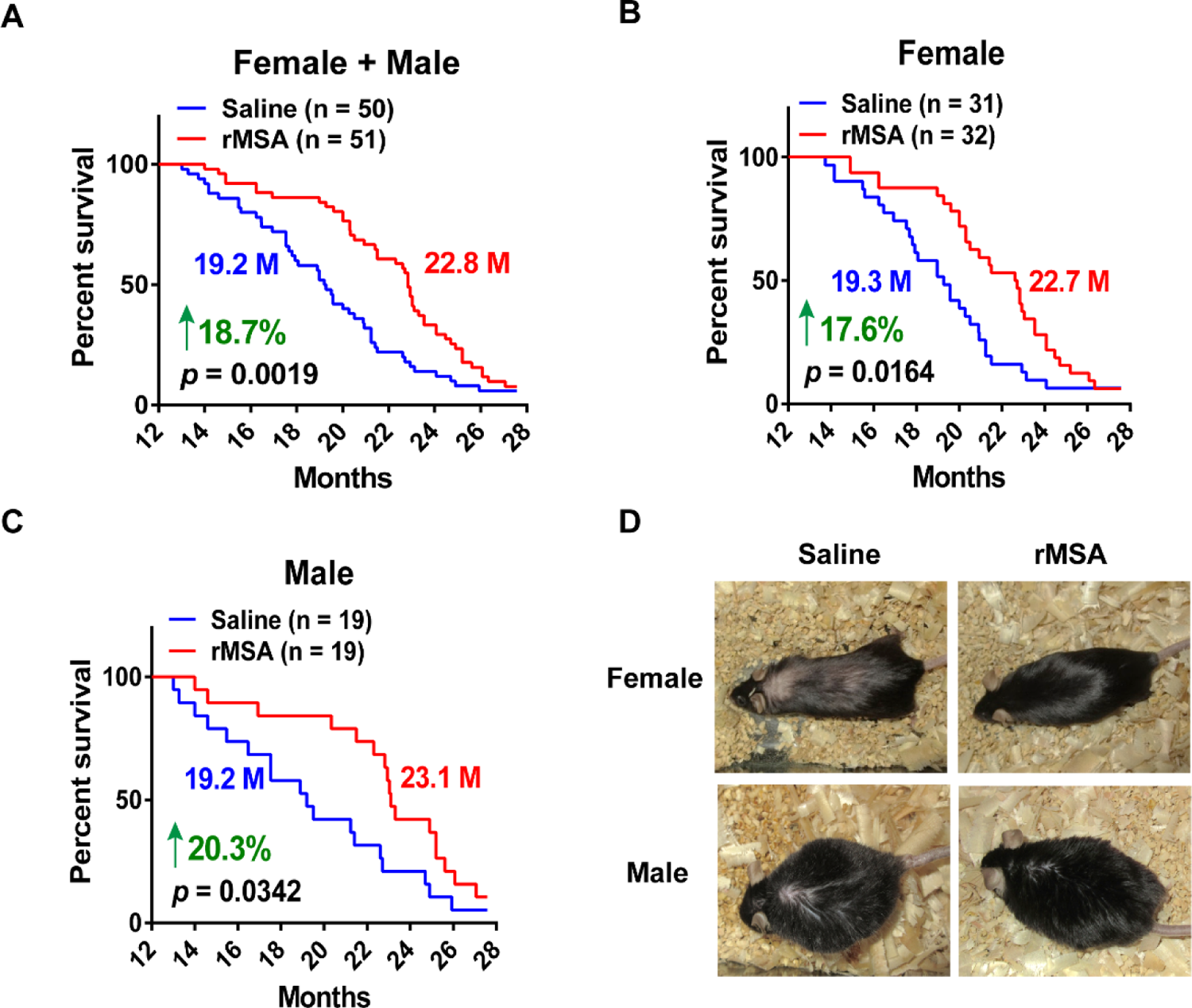
rMSA treatment increased the longevity in mice. (**A-C**) Survival curves of female (B) and male (C) mice treated with 1.5 mg rMSA per gram of body weight or isometric saline every 3 weeks. Median survivals (in months, M) and percentage increases are indicated. *p*-value was calculated by the log-rank (Mantel-Cox) test. n, number of mice used for each analysis. (**D**) Representative images of aged mice injected with rMSA or saline.

Moreover, qRT-PCR and blood biochemical analyses showed that both mRNA levels in the liver (Fig. S1A) and protein (Fig. S1B) levels in plasma of albumin underwent slight fluctuations before returning to normal within 8 days after the first injection. Major blood biochemical parameters remained constant in normal levels (Fig. S2A-N). In addition, to confirm whether the long-term treatment of saline or rMSA has various degrees of damages to organs, tissue sections of liver, kidney and heart were examined for any histopathological changes. Levels of α-SMA, a marker of myofibroblast activation in organ fibrosis [32], were measured in kidney (Fig. S3A-C), which showed no significant difference between saline- and rMSA-treated groups. To further verify the degree of renal fibrosis, Masson’s trichrome staining (Fig. S3D-F) and immunohistochemical staining of COL1A1 (Fig. S3G-I) were performed, which also showed no significant difference in kidneys of saline- and rMSA-treated mice. In the liver, α-SMA (Fig. S3J-L) and desmin (Fig. S3M-O) levels were measured to assess fibrosis levels, and no significantly differences were observed either. As for the heart, there was no significant difference in the collagen volume fraction of cardiac muscle by Masson’s trichrome staining (Fig. S3P-R). Furthermore, the lifespan of mice varies in different laboratories because of the different feeding conditions. The lifespans of the saline-treated mice in our study were similar to those of the unmanipulated wild type C57BL/6 mice in other studies [33, 34]. These phenomena suggest that rMSA treatment is safe for long-term use and can extend the lifespan of C57BL/6 mice.

### rMSA enhanced the function of skeletal muscle in mice

The elongation in mice lifespan triggered us to further explore whether the healthspan could also be improved. As the dysfunction in skeletal muscle was commonly observed during aging, we first detected the changes of grip strength in mice treated with rMSA or isometric saline for 8 months. rMSA-treated mice exhibited significantly increased forelimb grip strength from 177.9 g to 230.5 g (29.6% increased, *p* = 0.0002) in females and from 189.6 g to 222.5 g (17.4% increased, *p* = 0.0069) in males, as compared to saline-treated mice (Fig. 2A, B).

**Figure 2.**
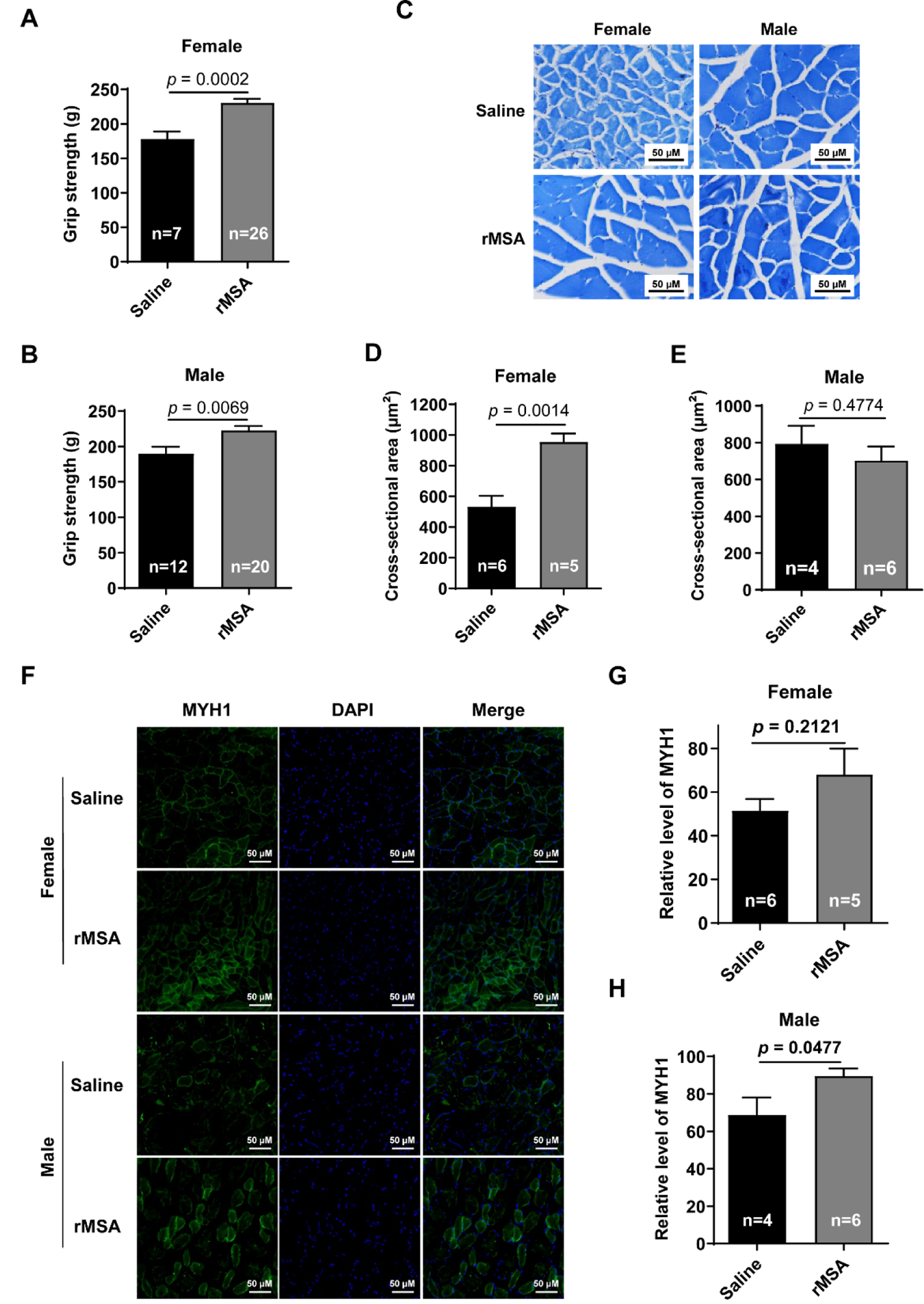
The effects of rMSA on the function of skeletal muscle in mice. (**A** and **B**) The grip strength of female (A) and male (B) mice. (**C**) The Toluidine Blue O staining of gastrocnemius muscle. Scale bar, 50 μm. (**D** and **E)** The cross-sectional area of myofibers in female (D) and male (E) mice. (**F**) The immunofluorescence staining for MYH1 (green) and DAPI (blue) in mice. Scale bar, 50 μm. (**G** and **H**) The relative level of MYH1 in female (G) and male (H) mice. Mice were treated with rMSA 1.5 mg per gram of body weight or isometric saline every 3 weeks for 8 months. All graphs represent mean with SEM, with *p* values calculated by the two-tail *t* test. n, number of mice used for each analysis

In order to evaluate the effect of rMSA injection on the *in vivo* skeletal muscle size and quality, we further performed histological analysis on gastrocnemius muscle (Fig. 2C). We found that the cross-sectional area of myofibers in rMSA-treated female mice were significantly increased (79.1% increased, *p* = 0.0014) than those in the saline group (Fig. 2D). However, similar phenomenon was not observed in male mice (Fig. 2E). We next investigated the expression level of slow MYH1 in rMSA and saline-treated group, another important parameter to evaluate the muscle strength (Fig. 2F). Male mice treated with rMSA presented significantly more slow MYH1 positive fibers than saline-treated mice (30.5% increased, *p* = 0.0477, Fig. 2H), while similar results were not obtained in female mice (Fig. 2G). Taken together, it was demonstrated that rMSA treatment enhanced the cross-sectional area of gastrocnemius fibers in female mice, and increased the level of slow MYH1 in male mice. We observed that rMSA had different effects on skeletal muscle of male and female mice, the variance of hormones and metabolic mechanisms may be one explanation for these differences. Interestingly, the improvement of skeletal muscle by rMSA treatment was coincident with that of the lifespan, demonstrating that rMSA most likely regulates both the lifespan and the healthspan based on the same fundamental principles.

### rMSA improved the spatial learning and memory of mice

We next investigated the effects of rMSA on aging-related impairment of memory using the Barnes Maze tests in male mice. rMSA-treated group exhibited a dramatic increase in the percentage of successful escape (73.2% *v.s.* 50.2%, 23.0% increased, *p* = 0.0016) compared to that of the saline-treated group (Fig. 3A, C). Meanwhile, the rMSA-treated male mice displayed significantly reduced primary escape latency (85.8 sec *v.s.* 133.4 sec, 47.6 sec faster, *p* < 0.0001) than the saline-treated mice (Fig. 3B, D). All these results demonstrated that rMSA treatment significantly improved the ability of spatial learning and memory in aging mice.

**Figure 3.**
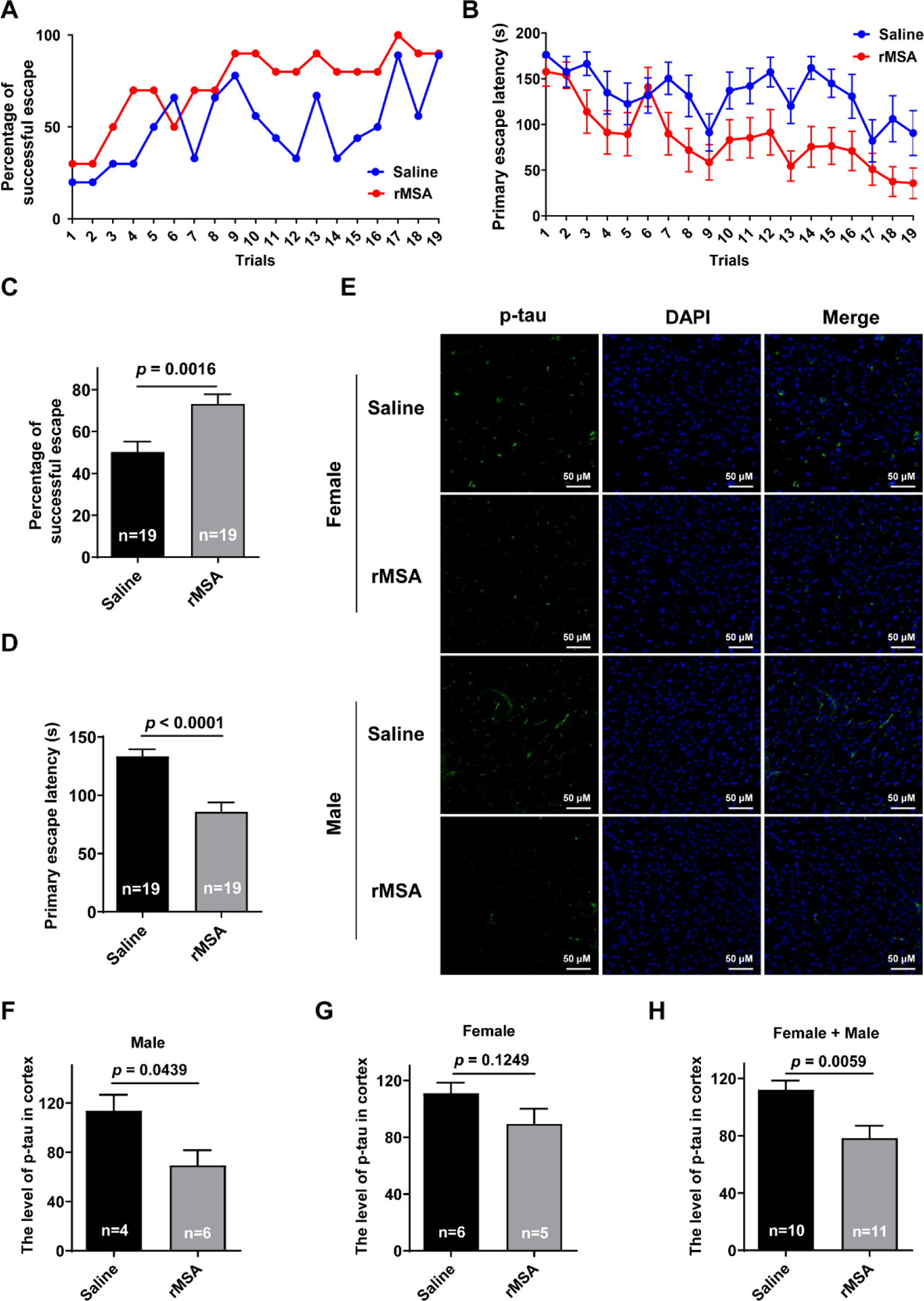
Effects of rMSA on the spatial learning and memory of mice. (**A** and **B**) Measurements of the percentage of successful escape (A) and the primary escape latency (B) of male mice. (**C**) The average of (A). n, the number of trials. (**D**) The average of (B). n, the number of trials. (**E**) The representative images of p-tau in the mice cortex. Scale bar, 50 μm. (**F-H**) The quantitative results of p-tau (green) and DAPI (blue) in male (F), female (G) and female + male (H) mice. n, the number of mice used for analysis. Mice were treated with rMSA 1.5 mg per gram of body weight or isometric saline every 3 weeks for 8 months. All graphs represent mean with SEM, with *p* values calculated by the two-tail *t* test

We then evaluated the histological changes associated with the memory using these groups of mice. Excitingly, the results of immunofluorescence staining in the cortex showed that the level of phosphorylated-tau (p-tau) was significantly decreased by rMSA treatment in male mice than that of the saline group (39.1% decreased, *p* = 0.0439, Fig. 3E, F). However, there was no significant discrepancy in female groups, though the level of p-tau in rMSA-treated mice was lower than that of the saline-treated mice (19.5% decreased, *p* = 0.1249, Fig. 3G). In sum, injection of rMSA decreased the p-tau level of mice (30.1% decreased, *p* = 0.0059, Fig. 3H), especially male mice. Furthermore, we suggest that effects of rMSA injection on memory improvement can relieve neurodegenerative disease during aging process..

### rMSA treatment improved four parameters related to aging

We proposed for the first time that the longevity can be enhanced by improving the status of free thiol, carbonyl, AGE, and Hcy which are the four parameters defining a young and undamaged protein. To verify this hypothesis, endogenous serum albumin samples of mice at 1.5-, 12-, and 28 months of age were purified respectively for comparison. During the aging process, serum albumin undergoes a series of changes in the four parameters: decreased level of free thiol and increased levels of carbonyl, AGE, and Hcy. The rMSA used in this study is even younger and less damaged than endogenous serum albumin from the young mice even at 1.5 months of age. The rMSA contains more free thiols (18.1 % increased, *p* = 0.0571) (Fig. 4A), equivalent level of carbonyl (Fig. 4B), less AGE (37.7% decreased, *p* = 0.0589) (Fig. 4C), and less Hcy (not detected in rMSA, *p* = 0.1215) (Fig. 4D). In addition, we need to emphasize here that no other damage was observed in our samples, because the molecular weight measured by mass spectrometry (Fig. S4A) is exactly the same as the theoretically calculated value [35]. In contrast, higher molecular weights were observed in endogenous albumin samples purified from mice serum at 1.5 months of age (Fig. S4B), which demonstrated more modifications on mouse serum albumin compared with rMSA. In sum, rMSA used in this study is not only “young” but also almost “undamaged”, which endows rMSA to offer more protection against unnecessary modifications and damages, and suggest that the four parameters could monitor the aging process. Here, “young” means that the rMSA is much fresher than the endogenous albumin from young mice at the age of only 1.5 months analyzed by the 4 parameters (free thiol, carbonyl, AGE, and Hcy). “Undamaged” theoretically means intact free thiol, no AGE, no carbonylation, and no homocysteinylation. In reality, due to the preparation process and detection methods, it is almost impossible to get such perfect sample.

**Figure 4.**
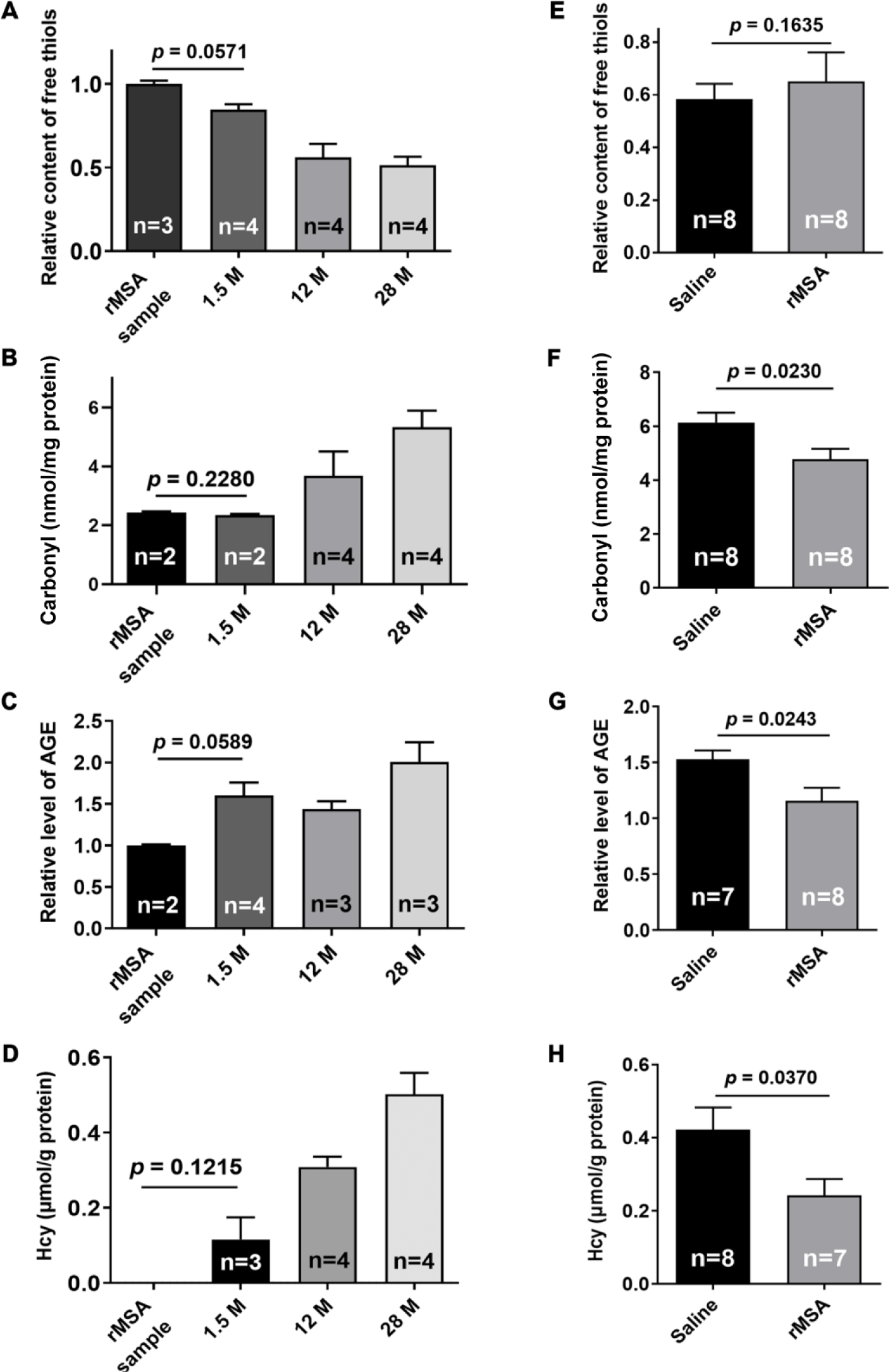
rMSA treatment improved four parameters related to aging. (**A-D**) The level of free thiol (A), carbonyl (B), AGE (C), and Hcy (D) of rMSA and endogenous albumin from serum samples of mice at 1.5-, 12-, and 28 months of age. (**E-H**) The level of free thiol (E), carbonyl (F), AGE (G), and Hcy (H) of endogenous albumin of mice treated with body weight-adjusted dosage of rMSA or isometric saline. All graphs represent mean with SEM, with *p* values calculated by the two-tail *t* test. n, number of mice used for each analysis.

In order to explore how young and undamaged rMSA improved the lifespan and healthspan of mice, 12-month-old mice were treated with 1.5 mg rMSA per gram of body weight or isometric saline every 3 weeks for 8 months. All serum samples were collected 21 days after the last injection. Compared with the saline-treated mice, the albumin from the rMSA-treated mice contained more free thiols (11.6% increased, *p* = 0.1635), much lower levels of carbonyl (22.1% decreased, *p* = 0.0230), AGE (24.4% decreased, *p* = 0.0243), and Hcy (42.6% decreased, *p* = 0.0370) (Fig. 4E-H). Taken together, young and undamaged rMSA provides a powerful protective function against oxidation of free thiol, carbonylation, AGE formation, and homocysteinylation.

## Discussion

Results showed that young and undamaged rMSA significantly improved the lifespan of mice with enhanced grip strength and memory. Our separate ongoing studies show that various physiological properties could be improved, such as immune responses, metabolic processes and cardiovascular functions. Further explorations will contribute to better understanding of the mechanism of young and undamaged rMSA on longevity.

Certainly, we realized that effects of rMSA and endogenous albumin on the longevity of mice should be compared in parallel. In order to perform this experiment, endogenous albumin should be prepared from mice at different ages ranging from very young to very old, whenever rMSA was used. However, endogenous mouse serum albumin of sufficient purity is not commercially available. Moreover, at least 20,000 mice at different ages were needed to purify sufficient amount of albumin at a purity greater than 99%, which is unethical.

A clinical trial whose purpose was to evaluate the beneficial effects of infusions of plasma from young donors (16-25 years old) to older adults (≥35 years old) was initiated in 2016 in the USA, but no result has been released so far (ClinicalTrials.gov Identifier: NCT02803554). Most recently, Conboy group reported rejuvenation of muscle, liver, and hippocampus of mice by exchanging old blood plasma with saline containing 5% endogenous albumin [36]. Pishel group reported that the injection of the plasma from young mice (2 to 4 months) cannot improve the median lifespan of middle-aged mice (10 to 12 months) [37]. Another clinical trial initiated by Grifols (ClinicalTrials.gov Identifier: NCT01561053, NCT00742417) showed that the plasma exchange with the replacement of human serum albumin significantly improved the cognitive performance in patients with Alzheimer’s disease compared with the control group [38]. However, neither of these studies used young and undamaged recombinant serum albumin, which makes the results not directly comparable.

In 2014, Wyss-Coray group reported that plasma from young mice can improve the learning and memory of old mice. Since albumin occupies about 50% of total plasma proteins, it most likely plays the most important role in this process, which was exactly what we found here. In order to achieve the maximal effect of rMSA on longevity, a variety of measures including optimal dosage, frequency, and drug delivery methods are being investigated. We predict that the concept of young and undamaged albumin increasing the longevity can also be applied to any other proteins such as immunoglobulins, fibrinogen, transferrin, transthyretin, and haptoglobin which are major plasma proteins.

It was well documented that the 4 parameters including free thiol, carbonyl, AGE, and Hcy are closely related to various diseases such as diabetes mellitus, cardiovascular diseases, adiposity, and Alzheimer’s disease [10, 23, 29, 39, 40, 41]. We discovered that longevity is intimately related to these four major parameters, based on which we defined the status of rMSA as “young and undamaged”. In addition, more parameters will be explored to enrich the definition of “young and undamaged” status in the future. It will be remarkable to see that a single young and undamaged protein (either recombinant or non-recombinant) HSA can increase the longevity of human beings, which will be initiated in the near future. If so, the combination of young and undamaged major plasma proteins can further increase the longevity. Ideally, all of the young and undamaged plasma proteins altogether can increase the longevity to the largest extent.

## Abbreviations

HSA: human serum albumin.
AGE: advanced glycation end-product.
Hcy: homocysteine.
rMSA: recombinant mouse serum albumin.
p-tau: phosphorylated microtubule-associated protein tau.
MYH1: myosin heavy chain I.
α-SMA: α-smooth muscle actin.
COL1A1: collagen I.
HCP: host cell proteins.
qRT-PCR: quantitative RT-PCR.
ELISA: enzyme-linked immunosorbent assay.
GAPDH: glyceraldehyde 3-phosphate dehydrogenase.
DAPI: 4′,6-diamidino-2-phenylindole.
DTNB: 5, 5’-Dithiobis-(2-nitrobenzoic acid).
Q-TOF: Quadrupole-Time of Flight.
K-S test: Kolmogorov-Smirnov test.

## Protein accession IDs (UniProtKB)

HSA: P02768

MSA: P07724

TAU: P10637

MYH1: Q5SX40

α-SMA: P62737

COL1A1: P11087

Desmin: P31001

## Author contributions

Yongzhang Luo, Yan Fu, Jiaze Tang, Anji Ju, Boya Li, Shaosen Zhang, and Yuanchao Gong designed the study; Yongzhang Luo, Yan Fu, Jiaze Tang, Anji Ju, and Boya Li wrote the manuscript, which was commented on by all authors; Jiaze Tang, Anji Ju, Boya Li, Shaosen Zhang, Yuanchao Gong, Boyuan Ma, Yi Jiang, and Hongyi Liu performed most of the experiments; Jiaze Tang, Anji Ju, Boya Li, Shaosen Zhang, Yuanchao Gong, Boyuan Ma, Yi Jiang and Hongyi Liu performed animal experiments; Jiaze Tang, Anji Ju, Boya Li, Shaosen Zhang, and Yuanchao Gong performed histological experiments; Jiaze Tang, Anji Ju, Boya Li, Yuanchao Gong, Shaosen Zhang, and Hongyi Liu performed biochemical experiments; Jiaze Tang, Anji Ju, Boya Li, Shaosen Zhang, Yuanchao Gong, Boyuan Ma, and Yi Jiang performed molecular experiments.

## Acknowledgments

We thank all the members of Shenzhen Protgen, Ltd. for kindly providing rMSA. Q-TOF-MS and additional technical assistance were performed by the Technology Center of Protein Research, Tsinghua University. We are also grateful to Laboratory Animal Research Center, Tsinghua University. We thank all the members in the Luo laboratory for their technical supports and insightful suggestions on this study.

## Supplemental Figure

**Figure S1.**
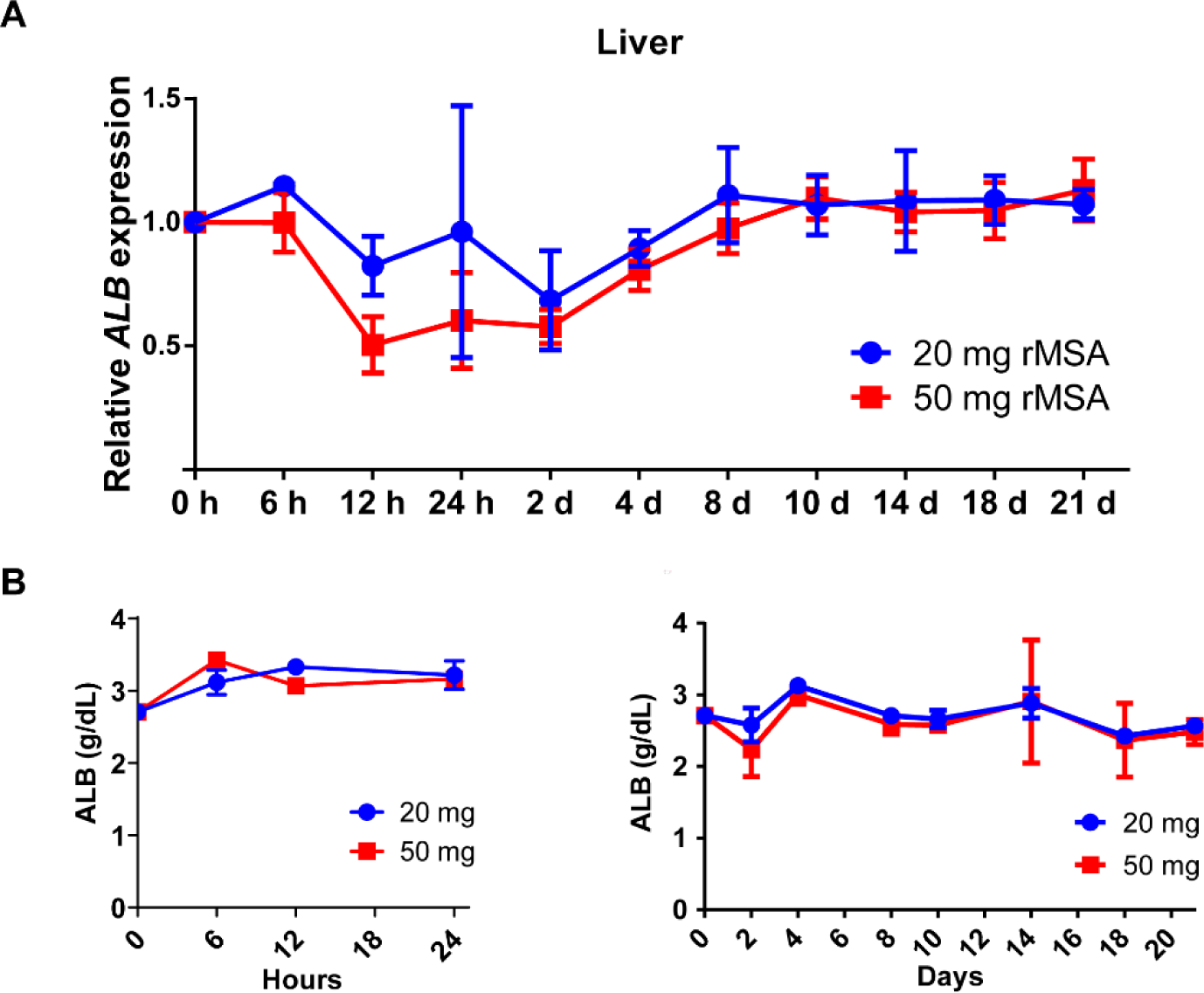
Effects of rMSA injection on the levels of albumin in mice (A) Dynamic expression levels of the albumin gene in the liver determined by qRT-PCR after the injection with 50 mg rMSA per mouse (n = 3). **(B)** Dynamic protein levels of the serum albumin within 1 day (left) and 2-21 days (right) after the injection with 20- or 50 mg rMSA per mouse (n = 3). All graphs represent mean with SEM.

**Figure S2.**
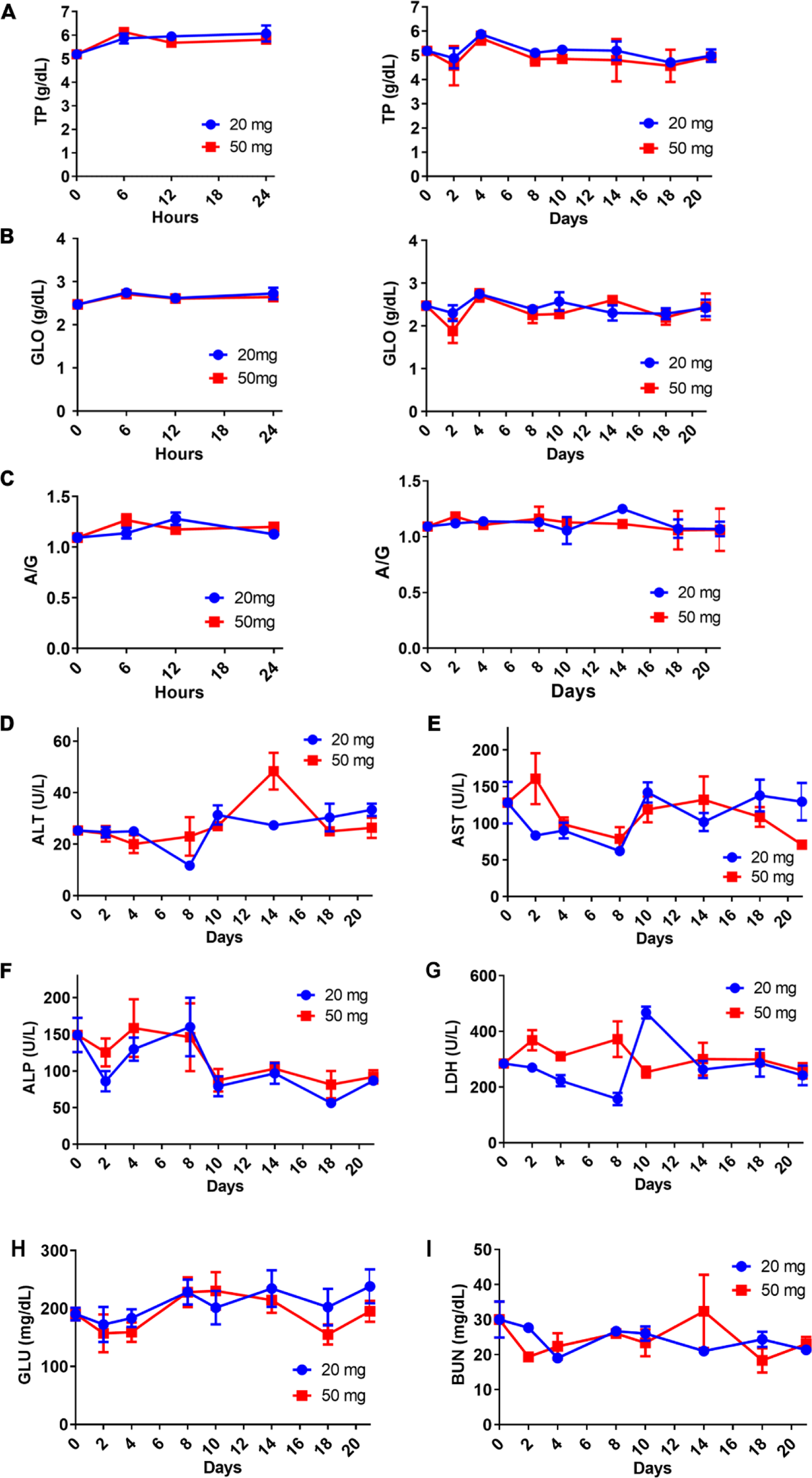
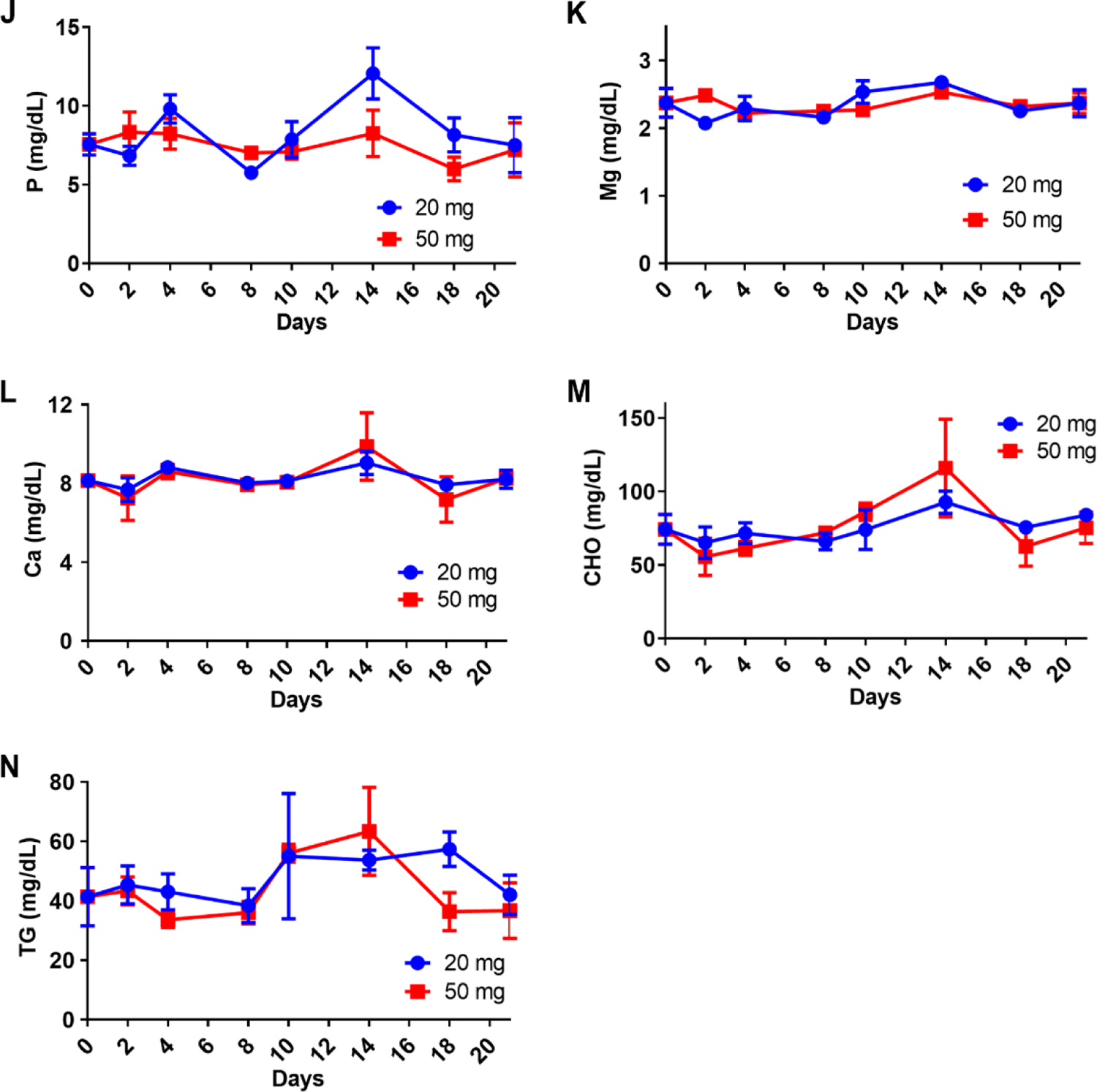
Effects of rMSA injection on the major blood biochemical parameters. (**A-C**) Dynamic total protein levels (A), total globulin levels (B), and the albumin/globulin ratio (C) within 1 day (left) and 2-21 days (right) after the injection with 20- or 50 mg rMSA per mouse (n = 3). (**D-N**) Dynamic levels of alanine transaminase (D), aspartate transaminase (E), alkaline phosphatase (F), lactate dehydrogenase (G), blood glucose (H), blood urea nitrogen (I), phosphorus (J), magnesium (K), calcium (L), cholesterol (M), and triglyceride (N) within 2-21 days after the injection with 20- or 50 mg rMSA per mouse (n = 3). All graphs represent mean with SEM.

**Figure S3.**
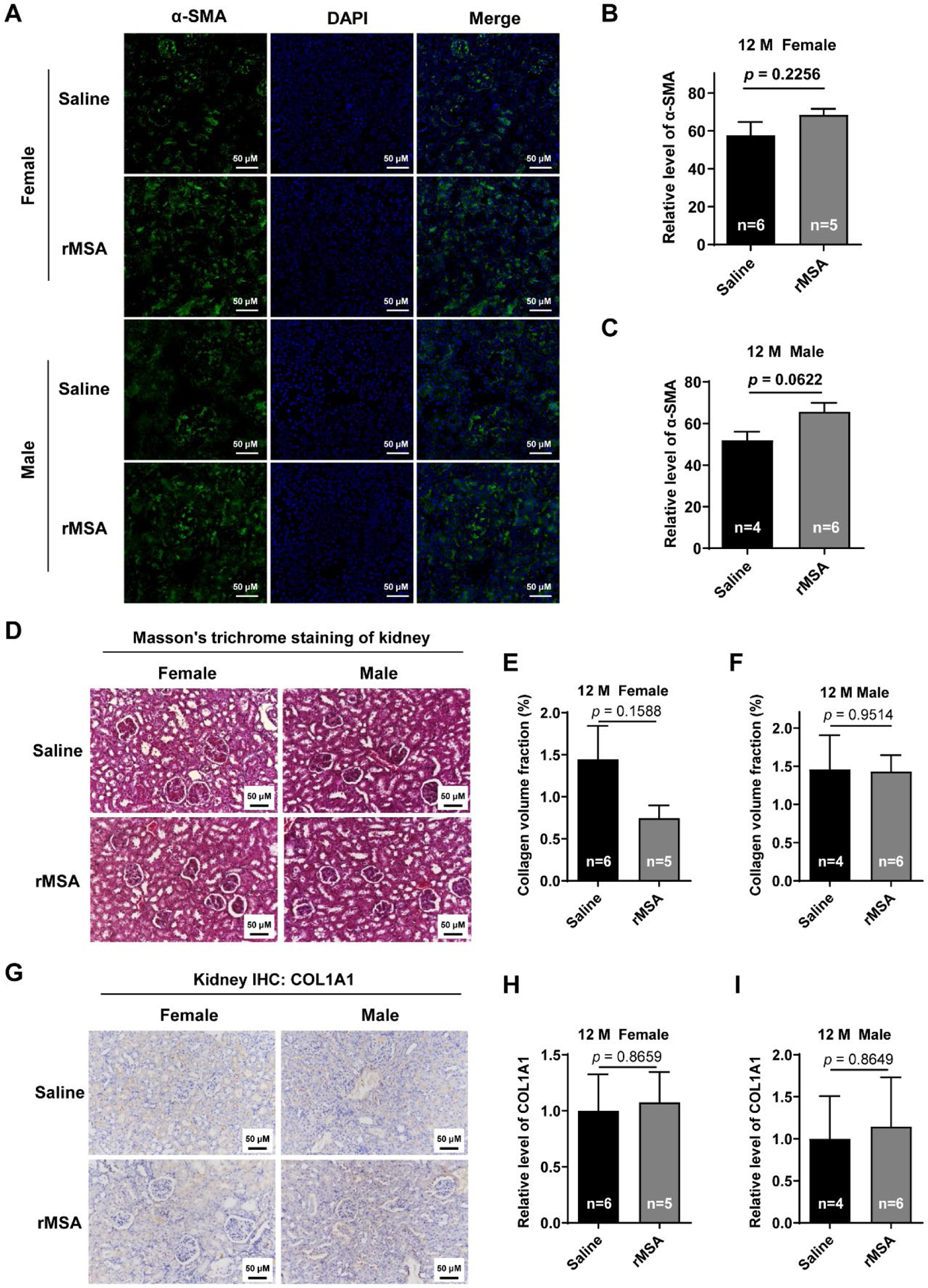
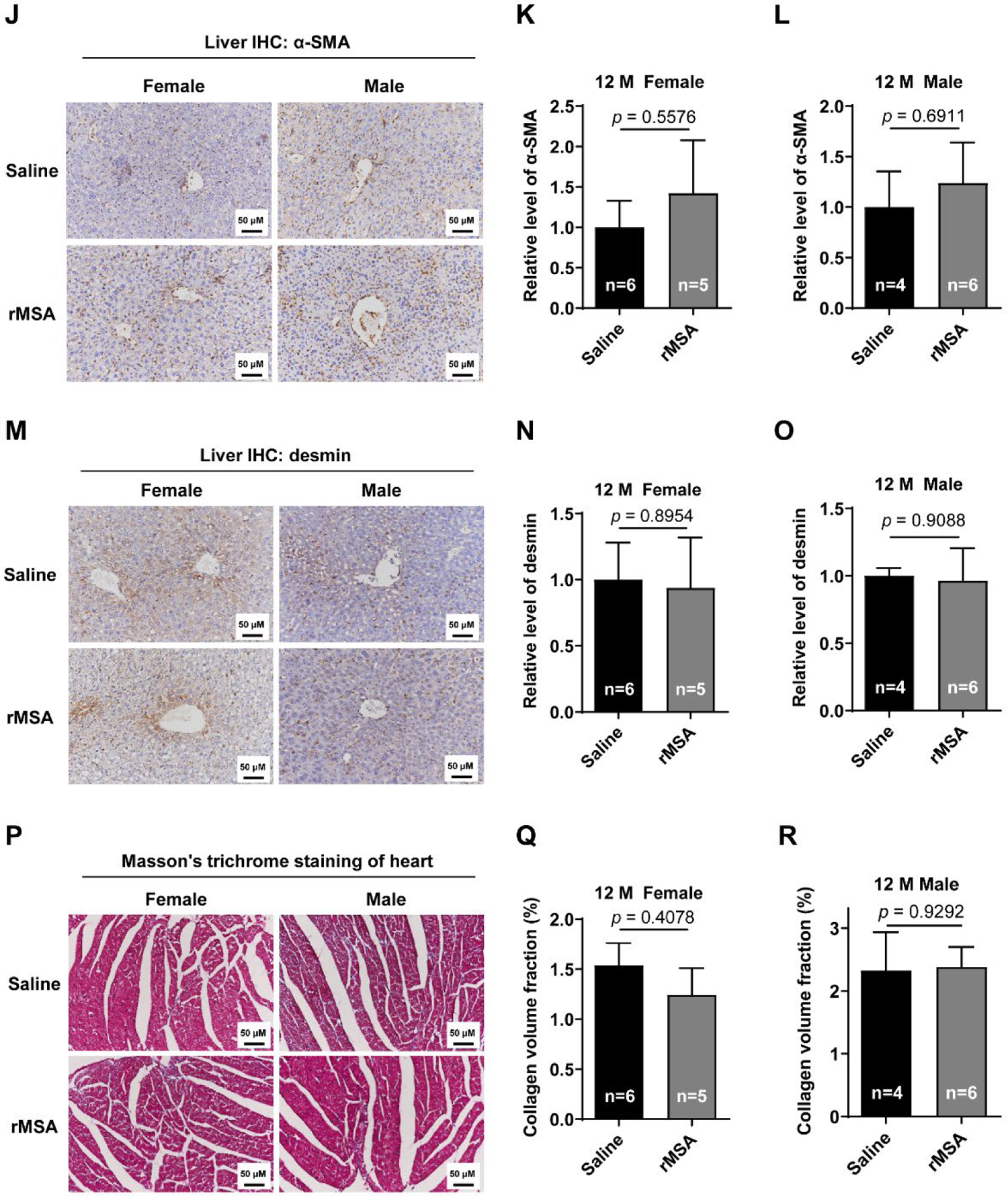
Effects of rMSA injection on the fibrosis level of kidney, liver and heart. (**A**) The representative images of α-SMA in mice kidney. Scale bar, 50 μm. (**B** and **C**) The quantitative results of α-SMA level in female (B) and male (C) mice. (**D**) The Masson’s trichrome staining of mice kidney. Scale bar, 50 μm. (**E** and **F)** The collagen volume fraction of the kidney in female (E) and male (F) mice. (**G**) The immunohistochemical staining for COL1A1 in mice kidney. Scale bar, 50 μm. (**H** and **I**) The relative level of COL1A1 in female (H) and male (I) mice. (**J**) The immunohistochemical staining for α-SMA in mice liver. Scale bar, 50 μm. (**K** and **L**) The relative level of α-SMA in female (K) and male (L) mice. (**M**) The immunohistochemical staining for desmin in mice liver. Scale bar, 50 μm. (**N** and **O**) The relative level of desmin in female (N) and male (O) mice. (**P**) The Masson’s trichrome staining of mice cardiac muscle. Scale bar, 50 μm. (**Q** and **R)** The collagen volume fraction of the cardiac muscle in female (Q) and male (R) mice. All graphs represent mean with SEM, with *p* values calculated by the two-tail *t* test. n, number of mice used for each analysis.

**Figure S4.**
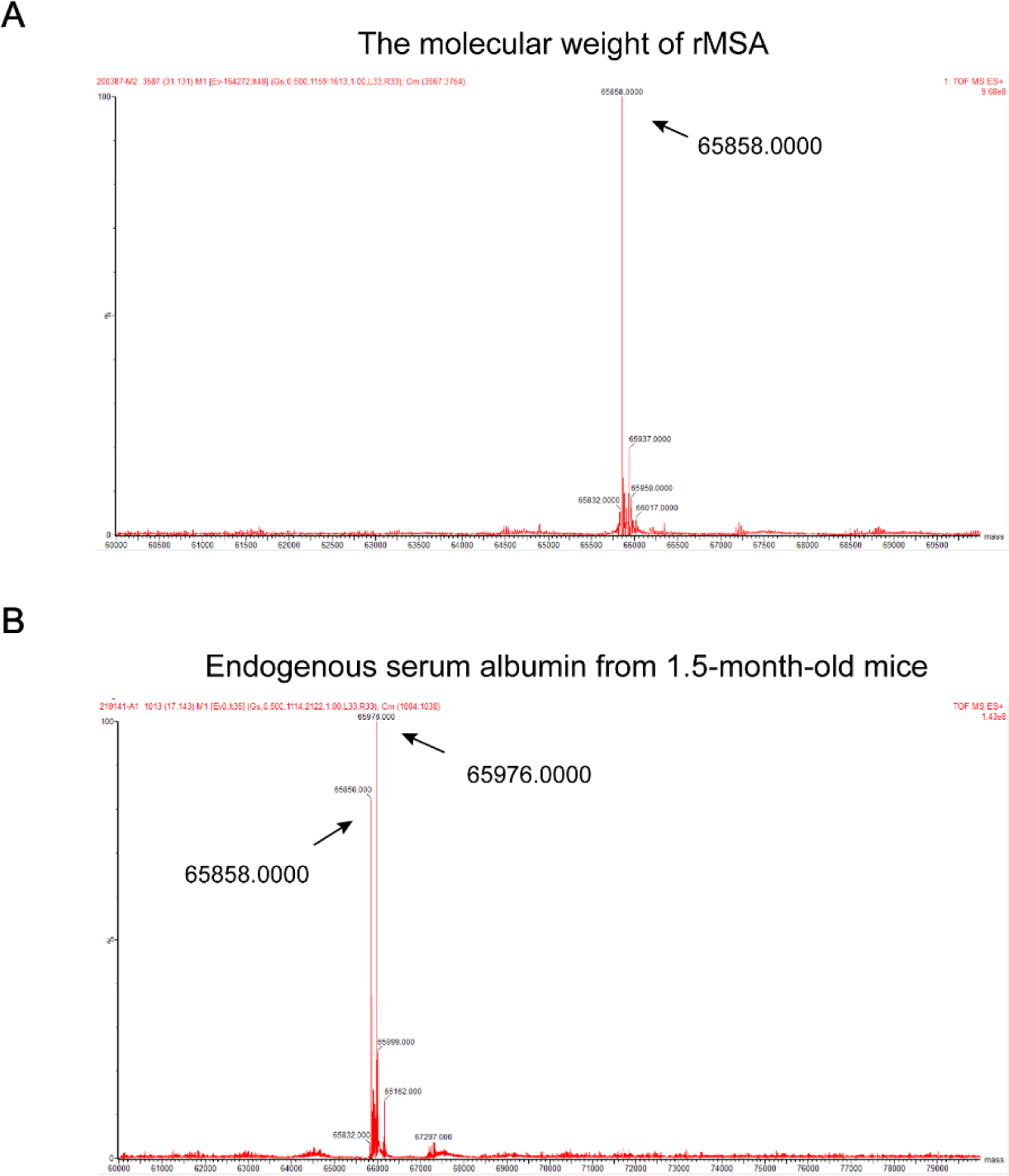
The mass spectrometry analysis of rMSA and endogenous mouse serum albumin. The molecular weight of rMSA (**A**) and endogenous albumin from serum sample of mice at 1.5 months of age (**B**) was verified *via* mass spectrometry (Q-TOF).

